# The Hypothalamic–Pituitary–Adrenal Axis Orchestrates Energy Homeostasis during Cold Exposure

**DOI:** 10.1101/2025.09.18.677113

**Authors:** Mireia Llerins Perez, Xinran Gao, Laura Heimerl, Yuanyuan Wang, Katharina Schnabl, Uwe Firzlaff, Harald Luksch, Martin Klingenspor, Nina Henriette Uhlenhaut, Dehua Wang, Yongguo Li

## Abstract

Cold exposure stimulates the sympathetic nervous system (SNS) to activate brown fat thermogenesis and maintain optimal body temperature, while simultaneously triggering compensatory hyperphagia to restore energy balance. The mechanisms coordinating energy expenditure and intake, however, remain unclear. Here, we reveal that the hypothalamic-pituitary-adrenal (HPA) axis plays a dual role in this process: endogenous adrenocorticotropic hormone (ACTH) directly stimulates the melanocortin-2 receptor (MC2R) in brown adipocytes to promote thermogenesis, whereas glucocorticoids drive cold-induced hyperphagia and act permissively to enhance ACTH-mediated energy expenditure. These findings uncover previously unrecognized functions of the HPA axis and a delicate hormonal interplay that orchestrates energy homeostasis during cold stress. Targeting these pathways may offer novel strategies to mitigate hyperphagic responses associated with increased energy expenditure, with potential implications for obesity treatment.

**Graphical abstract:** 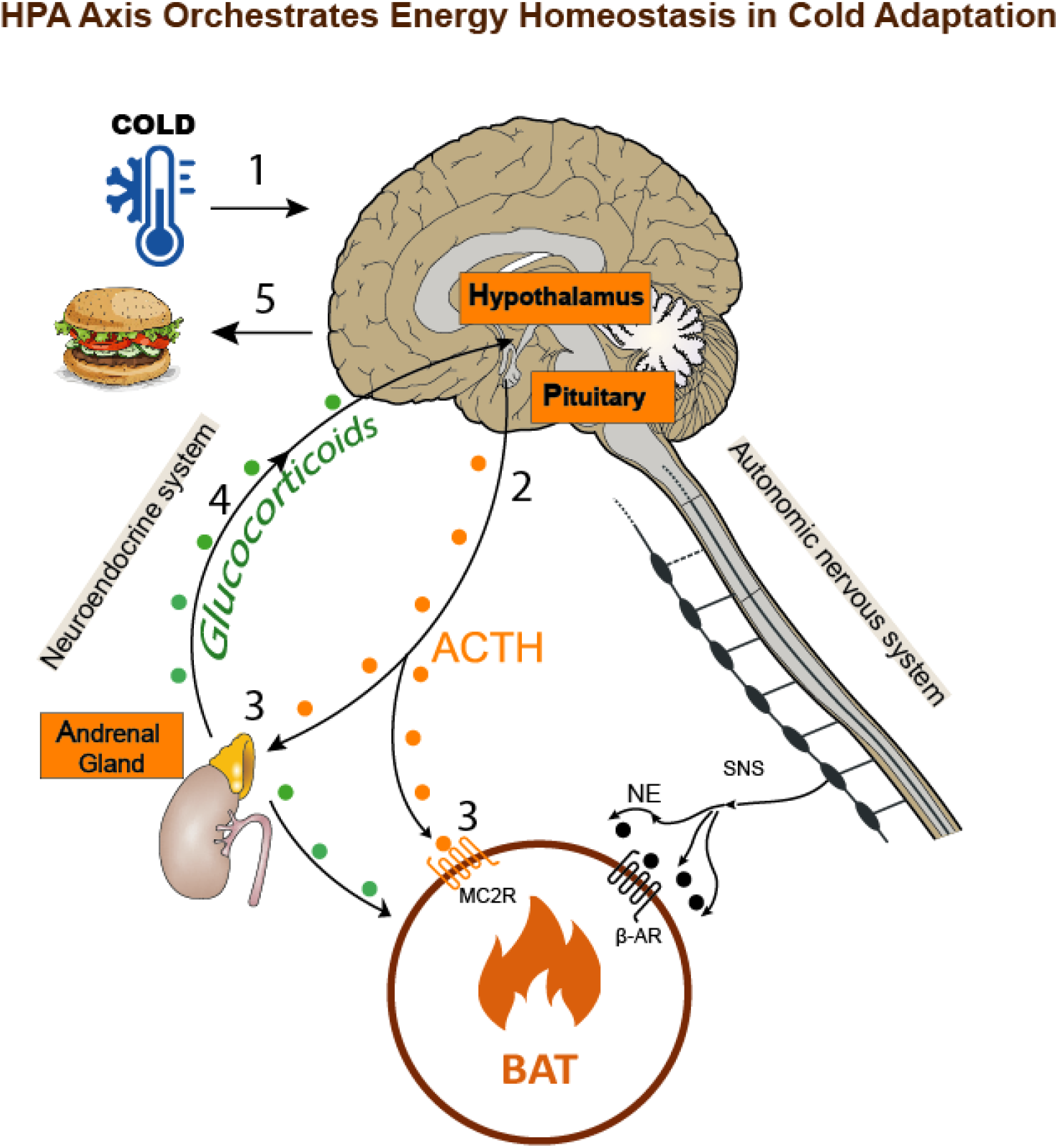

**Highlights:**

- The HPA axis orchestrates cold adaptation.
- ACTH activates brown fat thermogenesis *in vitro* and *in vivo* via activating MC2R.
- Glucocorticoids drive cold-induced hyperphagia.
- Glucocorticoids have a permissive effect on ACTH function in brown adipocytes.

## Introduction

Due to its high energy cost, thermogenesis in mammals is tightly controlled and exquisitely coordinated by multiple hormones and organ systems to maintain whole-body energy homeostasis (Silva, 2006). Upon cold exposure, signals evoked by increased heat loss and thermoreception via cutaneous thermoreceptors converge to increase heat generation by activating the sympathetic nervous system (SNS) (Morrison & Nakamura, 2011; Tan & Knight, 2018). In small mammals, brown adipose tissue (BAT) is the main heater organ for cold-induced thermogenesis (Klingenspor, 2003; Li, Lasar, et al., 2014). Catecholamines released from sympathetic nerve endings in BAT act via β3-adrenoceptors to stimulate lipolysis and activate the uncoupling protein-1 (UCP-1), which uncouples oxidative phosphorylation to produce heat via oxidation of fatty acids and glucose (Cannon & Nedergaard, 2004). Energy combustion would lead to an energy deficit without compensation. To maintain energy homeostasis during cold exposure, the increased energy demands of thermogenesis must be counterbalanced by increased energy intake, known as cold-induced hyperphagia (Cannon & Nedergaard, 2009; Deem et al., 2020a; Lal et al., 2023; Melnyk & Himms-Hagen, 1998; Yang et al., 2021). As such, thermoregulation and feeding are fundamental intertwined aspects of mammalian biology (Brobeck, 1948; Li et al., 2018). Nevertheless, how energy expenditure and energy intake are coordinated to maintain energy homeostasis remains poorly understood. Since energy compensation is a major hurdle for obesity treatment (Doucet et al., 2018), a more thorough understanding of the processes that coordinate thermogenesis and feeding could have important therapeutic implications for obesity-related metabolic disease.

Notably, cold exposure not only activates the SNS but also stimulates the hypothalamic-pituitary-adrenal axis (HPA), which is a major neuroendocrine signalling system involved in physiological homeostasis and the stress response (McCarthy et al., 1993). Upon exposure to cold stress, neurons in the paraventricular nucleus of the hypothalamus secrete corticotropin-releasing hormone (CRH) from nerve terminals in the median eminence into the hypothalamo-hypophyseal portal circulation, which stimulates the production and release of adrenocorticotropic hormone (ACTH) from the anterior pituitary (Axelrod & Reisine, 1984). ACTH, in turn, stimulates the release of glucocorticoids from the adrenal cortex. Circulating glucocorticoids, the end-products of the HPA axis, exert their pleiotropic effects in almost all tissues and organs and play a fundamental role in facilitating adaptation to stress and restoring homeostasis through integrating permissive, suppressive, stimulatory, and/or anticipatory actions (Sapolsky et al., 2000). It is thus generally believed that the primary function of the activated HPA axis during cold exposure is to manage cold stress. Since cold exposure enhances ACTH and subsequently corticosterone release, it is very likely that these hormones, even in a synergystic manner, play a role in cold-induced thermogenesis and energy homeostasis, which has not been rigorously tested so far.

Consistent with the above mentioned scenario, supraphysiological doses of ACTH can activate murine brown adipocyte thermogenesis *in vitro* (Schnabl et al., 2018) and glucose uptake *in vivo* (van den Beukel et al., 2014). ACTH signaling depends on the ACTH receptor complex, which is composed of the melanocortin 2 receptor (MC2R), a Gs protein-coupled receptor (GPCR), and the MC2R accessory protein (MRAP), which is essential not only for receptor responsiveness to ACTH but also for the transport of MC2R to the cell surface(Cooray et al., 2008; Metherell et al., 2005). Binding of ACTH to MC2R in the presence of MRAP leads to cAMP/PKA activation and lipolysis stimulation (Lefkowitz et al., 1970). It is thus not that surprising to observe a thermogenic effect when using supraphysiological levels of ACTH, given that lipolysis stimulation in brown fat typically leads to activation of UCP1-medidated thermogenesis (Braun et al., 2018; Li et al., 2022). Nevertheless, it remains to be ascertained whether the physiological rise in ACTH levels *in vivo* is sufficient to activate BAT thermogenesis and contribute to acute cold adaptation. In addition, it is still not fully understood how MC2R and its downstream signaling pathway control thermogenesis in brown adipose tissue.

ACTH-stimulated glucocorticoids (GCs) exert widespread effects, playing a central role in maintaining homeostasis and orchestrating adaptive responses to stress (Munck et al., 1984). Beyond limiting excessive stress-induced responses, GCs act permissively, enhancing the actions of other hormones without directly eliciting those effects themselves (Sapolsky et al., 2000). Notably, GCs enable catecholamines to mobilize energy (lipolysis), maintain vascular tone (vasoconstriction), and regulate blood pressure (cardiac output) (Hadcock & Malbon, 1988a; Kalsner, 1969; Ullian, 1999). Despite these well-established roles, potential additional functions of glucocorticoids during cold exposure remain largely unexplored, representing an important knowledge gap in our understanding of stress and metabolic regulation.

In this study, we rigorously investigated the physiological role of the HPA axis in brown fat metabolism and energy homeostasis during cold exposure. Using gain- and loss-of-function approaches to manipulate HPA activity, we found that exogenous ACTH acutely increased energy expenditure to a level comparable to norepinephrine (NE), while blockade of endogenous ACTH using a neutralizing antibody or the MC2R antagonist CRN04894 impaired cold-induced thermogenesis. Interestingly, inhibition of ACTH signaling reduced food intake during cold exposure, whereas infusion of dexamethasone, a potent synthetic glucocorticoid, further enhanced feeding. These findings indicate that the HPA axis not only drives cold-induced heat production but also mediates compensatory energy intake to support increased energy expenditure. Furthermore, glucocorticoids act permissively to support ACTH-driven responses. Collectively, our results reveal previously unrecognized functions of the HPA axis in coordinating energy balance during cold exposure, highlighting its dual role in regulating cold-induced energy expenditure and intake to maintain energy balance.

## Methods

### Animals

Male C57BL/6J and male 129S1/SvImJ mice (UCP1-KO mice and wild-type littermates) were used for the *in vivo* experiments. Animal experiments were approved by the German animal welfare authorities at the district government (approval no. 55.2-1-54-2532-34-2016) and by the Animal Experimental Ethics Committee of School of Life Sciences, Shandong University (approval no. SYDWLL-2021-96). Mice were bred at the specific-pathogen free animal facility. Animals had *ad libitum* access to food and water and were maintained at 22 ± 1°C and 50%–60% relative humidity in a 12 h:12 h light:dark cycle. Male mice, aged 5 to 6 weeks were used for primary cultures of brown adipocytes.

### Cell culture

Interscapular brown adipose tissue was isolated from 5-week-old 129S1/SvImJ mice (UCP1-knockout and wild-type littermates; n = 5 per experiment, pooled) and digested with collagenase as previously described (Li, Bolze, et al., 2014; Oeckl et al., 2020). Stromal vascular fraction (SVF) cells were plated and cultured to confluence or immortalized using a retrovirus-mediated expression of the simian virus 40 large T antigen. Differentiation was initiated for 2 days using adipocyte culture medium (DMEM with 10% heat-inactivated FBS, penicillin/streptomycin, and gentamicin) supplemented with 5 μg/mL insulin, 1 nM 3,3′,5-triiodo-L-thyronine (T3), 125 μM indomethacin, 0.5 mM isobutylmethylxanthine (IBMX), 1 μM dexamethasone and 1 µM rosiglitazone. Cells were then maintained for 6 additional days in adipocyte differentiation medium containing 5 μg/mL insulin, 1 nM T3, and 1 µM rosiglitazone, with media being replaced every 2 days. Fully differentiated adipocytes with prominent lipid droplet accumulation were used for downstream experiments.

### Respiration assays

The cellular oxygen consumption rate (OCR) of primary and immortalized adipocytes was determined using a Seahorse XF96 Pro Analyzer (Agilent technologies) as described previously (Li, Fromme, et al., 2014; Oeckl et al., 2020). Briefly, primary SVFs or immortalized cells or CIRSPR-Cas9 based gene KO cells were cultured and differentiated in XF96 microplates. After differentiation, cells were washed with prewarmed basal assay medium [DMEM with 25 mM glucose, 31 mM NaCl, 2 mM GlutaMax, pH 7.4], then incubated for 1 h at 37°C in basal assay medium containing 1%–2% fatty acid–free BSA. 10× chemical reagents in basal assay medium (without BSA) were loaded into sensor cartridge ports. Basal respiration was first measured, followed by oligomycin (Sigma-Aldrich) (5 μM) to inhibit coupled respiration, ACTH (Bachem) (0.25 µM) to assess UCP1-dependent uncoupling, FCCP (Sigma-Aldrich) (7.5 μM) to measure maximal capacity, and antimycin A (Sigma-Aldrich) (5 μM) to determine non-mitochondrial respiration. In some experiments, cells were pretreated with the ACTH-neutralizing antibody ALD1613 (AntibodySystem) at a 1:2 molar ratio of ACTH to antibody or the MC2R antagonist CRN04894 (MedChemExpress) with varying doses ranging from 0.1-100 µM. Oxygen consumption rates were calculated using prism software. Experiments were repeated at least three times with similar results.

### CRISPR-Cas9 based loss-of-function assay

For electroporation-based CRISPR/Cas9 gene knockout, the Neon Transfection System (10 μL Kit, Thermo Fisher, #MPK10096) was used to deliver the assembled Cas9 ribonucleoprotein (RNP) complex to preadipocytes. The RNP complex, containing target-specific guide RNA (annealed crRNA:TracrRNA duplex, 1.8 μM) and Alt-R® S.p. Cas9 Nuclease V3 (1.5 μM, all IDT), was assembled prior to use. After washing with PBS to remove the serum, cell pellets were resuspended in Neon Resuspension Buffer R to a final cell density of 0.5-6x10^6^ cells per electroporation (in 10 μL). Cell suspensions were mixed with the RNP complex and Cas9 electroporation enhancer (1.8 μM, IDT). Electroporation was performed using the Neon Transfection System with one pulse at 1750 V and 20 ms width. Cells were subsequently cultured in antibiotic-free culture medium, were let to recover overnight, and were then expanded and differentiated for downstream assays. The *Mc2r* gene was targeted with crRNAs 5′ ACAATCGGAGTTATTTCTTG3′ and 5′ TCATCACCCTAACAATTATC3′. *The Mrap* gene was targeted with crRNA 5′ CACCAGCTATGAGTATTAC3’ and 5′ GGAGCACCACGAAGGTAGCC′.

### Indirect calorimetry

Indirect calorimetry was conducted using an open respirometry system (LabMaster, TSE Systems, Germany) as described previously (Li et al., 2018; Wang et al., 2021). For measuring NE or ACTH-induced energy expenditure, mice were placed in 1.5 L metabolic cages without food and water, kept at a climate cabinet preconditioned to 30 °C and connected to the indirect calorimetry setup (Phenomaster, TSE Systems, Germany) to measure basal metabolic rate (BMR) during fasting. The air from the cages was extracted over a period of 1 min every 3 min with a flow rate of 24 l/h, dried in a cooling trap and analyzed for O_2_ content. O_2_ consumption [ml/h] was calculated via comparison of the air from the cages with the air from an empty reference cage. The lowest mean of three consecutive oxygen consumption values was calculated as BMR. After BMR measurements, mice were i.p. injected either PBS, NE (1 mg/kg) or ACTH (1 mg/kg) and the metabolic rate was measured in the calorimetry chamber for about one hour at 27°C to avoid hyperthermia. For cold-induced energy expenditure, O_2_ consumption was recorded using the same TSE setting.

Food intake (FI) was monitored using an automated monitoring system (TSE LabMaster Home Cage Activity, Bad Homburg, Germany). The food baskets were connected to high precision balances to record FI in 1-min bins.

For the endogenous ACTH blocking experiment, mice were i.p. injected with either vehicle or 5 µg/mouse of the anti-ACTH antibody ALD1613 (AntibodySystem) 1 hour prior to measuring energy expenditure upon cold exposure. For the MC2R antagonism experiment, 60 mg/kg of the MC2R antagonist CRN04894 (MedChemExpress) were administered via oral gavage 1 hour before measuring energy expenditure upon cold exposure.

### Monitoring of body temperature and iBAT temperature

Mice were anesthetized by combined i.p. injection of medetomidine (0.5 mg/kg), midazolam (5 mg/kg) and fentanyl (0.05 mg/kg). The interscapular region was opened by a small incision and the temperature probe was placed to the interscapular BAT pad. Meanwhile, a 2nd temperature probe was inserted into the colon (2 cm deep) to measure core body temperature. ACTH was injected directly into the BAT lobe at specified doses.

### Gene expression analysis (qRT–PCR)

RNA was extracted from cultured cells using TRIsure, purified with SV Total RNA Isolation System, Promega and reverse transcribed using SensiFAST cDNA Synthesis Kit (BIOLINE). The resultant cDNA was analyzed by qRT–PCR. Briefly, 1 µL of 1:10 diuluted cDNA and 400 nmol of each primer were mixed with SensiMix SYBR Master Mix No-ROX (Bioline) into a 10 µL reaction volume. Reactions were performed in 384-well format using the QuantStudio™ 5 Real-Time PCR System (ThermoFisher). The RNA abundance of each gene was normalized to the expression of the housekeeping control Hprt, and the 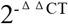 method was applied to calculate changes in gene expression. The following primers were used: Mc2r Forward: CAGTGCTCACCTTCACATCG; Reverse: AAGGATGGTTAGTGTCATGGC; Mrap Forward: TGTGGTTGAGCCTGGCTACCTT; Reverse: GGAGGTTGAAGCTGTGAGTCCA; Hprt Forward: TCAGTCAACGGGGGACATAAA; Reverse: GGGGCTGTACTGCTTAACCAG.

### Measurement of plasma ACTH and Glucocorticoid concentrations

Blood was collected at the time the mice were killed. Plasma ACTH and corticosterone levels were determined by ELISA using the kit purchased from Shanghai Lianshuo Biotech Co., Ltd. (AE91004Mu, AE90708Mu, AMEKO, Shanghai, China) following the manufacturer’s instructions. Concentrations were calculated using a standard curve generated with ACTH and corticosterone standards included in the kit.

### Luciferase-based GloSensor cAMP assay

cAMP production was assessed in differentiated immortalized brown preadipocytes transduced with a lentiviral pCDH-PGK-pGloSensor-22F reporter using the luminescence-based GloSensor™ cAMP assay (Promega). Cells were cultured and differentiated in 96-well white plates (Corning, 3610). On day 7–8 of differentiation, cells were either pretreated (PBS wash, with or without 1 μM dexamethasone for 30 min, followed by with or without 0.25 μM ACTH for 1 h, then returned to differentiation medium for ∼22 h) or left untreated as controls.

Prior to measurement, cells were equilibrated in assay medium (88% CO_2_-independent medium, 10% FBS, 2% GloSensor reagent) for 1 h at 37 °C and 30 min at room temperature. The assay medium was then supplemented with 10 µl/well of a solution of IBMX (Sigma-Aldrich, I5879, final concentration: 200 µM), a non-selective phosphodiesterase inhibitor that helps stabilizing the cAMP signal by preventing its degradation, and incubation at room temperature in the dark for 30 min.

Luminescence intensity was measured over time using a luminometer (BioTek Synergy H1, Agilent Technologies). Measurements were obtained at 2.5-minute intervals with an integration time of 1 s. Luminescence was first assessed for 10 minutes in a kinetic cycle to establish baseline luminescence values. This was followed by the addition of a 10 µl/well of a solution of ACTH in CO_2_-independent medium with a final concentration of 0.25 µM per well. After 1 s of orbital shaking, luminescence was measured in a kinetic cycle for 1 hour.

### Lipolysis assay

Differentiated adipocytes were washed with PBS and then incubated in lipolysis medium containing either vehicle, 1 µM isoproterenol (Sigma-Aldrich, I6504) or 0.25 µM ACTH (Bachem, 4034845). The lipolysis medium consisted of DMEM (Gibco, 11966025), 1 g/l glucose (Carl Roth, X997.2), and 2% fatty acid free BSA (Sigma-Aldrich, A3803). Cells were incubated at 37°C and 5% CO_2_ for four hours. Cell culture medium was collected and frozen at -20ºC. To measure glycerol content, 15 µl of the media samples were incubated for 15 min with 75 µl of free glycerol reagent (Sigma-Aldrich, F6428) at room temperature. Absorption was measured at 540 nm with a multiplate reader (PerkinElmer, EnSpire). Glycerol release was calculated using a glycerol standard (Sigma-Aldrich, G7793).

### Diet-induced obese mice study

Diet-induced obese (DIO) male C57BL6/L mice 17 to 18 weeks old, maintained on a high fat diet were ordered from the Jinan Pengyue Experimental Animal Breeding Center (Jinan, China) and were used in the study. Mice were individually housed in a temperature-controlled (24°C) facility with 12 hour light/dark cycle and free access to high fat diet (HFD) (D12492) and water. After a minimum of 2 weeks acclimation to the facility, the mice were randomized according to their body weight, so each experimental group of animals would have similar body weight. Mice were then daily injected with CL316,243 (CL; 1.0 mg/kg i.p.), ACTH (1 mg/kg i.p.) or saline for 6 consecutive days while feeding with a HFD. Daily body mass changes and metabolic rates were monitored during the treatment.

### Quantification and Statistical Analysis

Statistical analysis was performed using GraphPad Prism (GraphPad Software). Data are presented as the mean ± SD. Two-tailed Student’s t tests were used for single comparisons and analysis of variance (ANOVA) with Tukey’s post hoc tests for multiple comparisons. For DIO body-weight change, ANCOVA was used with baseline body weight as a covariate to control for initial differences. P values below 0.05 were considered significant.

## Results

### Exogenous ACTH treatment boosts energy expenditure *in vitro*,and *in vivo*

It has been extensively reported that ACTH can stimulate lipolysis in rodent adipose tissues and mouse adipocytes via MC2R-dependent cAMP/PKA activation (Boston, 1999; Boston & Cone, 1996; Cho et al., 2005; Hollenberg et al., 1961; White & Engel, 1958). As lipolysis stimulation in brown fat typically leads to activation of UCP1-mediated thermogenesis, the thermogenic potential of ACTH has been a long-standing interest (Schnabl et al., 2018; van den Beukel et al., 2014). However, whether ACTH-stimulated energy expenditure is mediated by UCP1 and the physiological importance of this thermogenic effect remain to be determined. To fill this knowledge gap, we first stringently tested the acute direct thermogenic effects of ACTH in multiple conditions, such as *in vitro* and *in vivo*. In line with our previous finding, ACTH stimulated the oxygen consumption rate of wild-type but not UCP1 knockout (UCP1KO) brown adipocytes (Fig.1A). As shown in Fig. 1B, the ACTH receptor melanocortin 2 receptor (MC2R) and its functional component MC2R-associated protein (MRAP) are expressed in adipocytes during differentiation, indicating a direct stimulatory effect of ACTH on brown adipocytes. Consistently, local injection of ACTH into the interscapular brown fat lobe in anesthetized mice increased BAT temperature and consequentially raised body temperature (Fig.1C). Lastly, we performed indirect calorimetry measurement of energy expenditure in living animals upon acute ACTH injection. We show that an exogenous ACTH injection acutely increases energy expenditure compared to the PBS control (Fig.1D).

**Figure 1.**
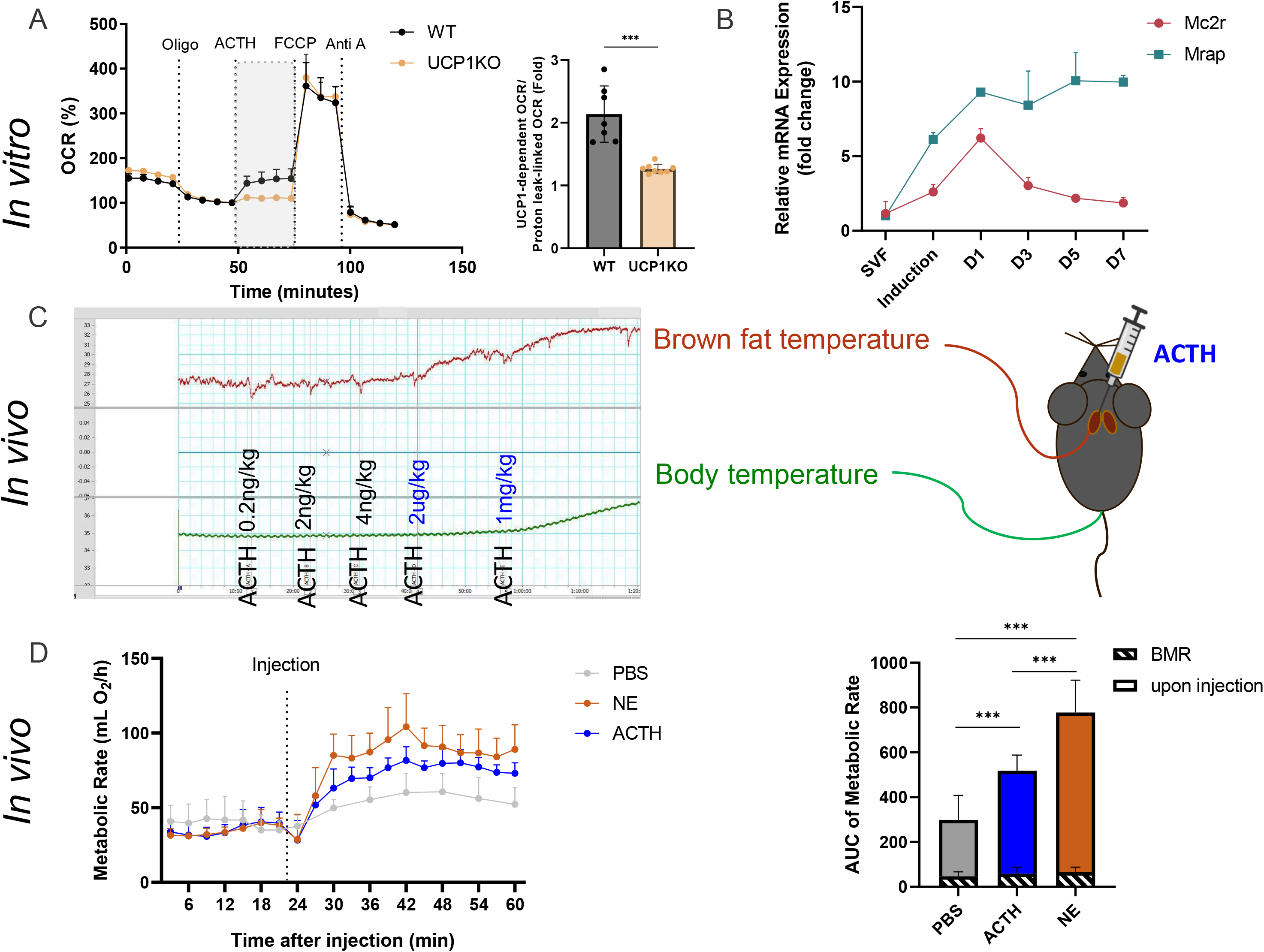
Exogenous ACTH activated brown fat thermogenesis under *in vitro, ex vivo* and *in vivo* conditions. (A) Time course of oxygen consumption rate of UCP1 WT and knockout (KO) brown adipocytes recorded by microplate-based respirometry (Seahorse XF96 Analyzer) under basal conditions and during successive addition of 5 µM oligomycin (oligo), 0.25 µM ACTH, 7.5 µM FCCP, and 5 µM antimycin A (Anti A). Data are expressed as percentage of basal leak respiration.UCP1-mediated uncoupled respiration (highlighted in grey) is expressed as fold increase of basal leak respiration after stimulation with ACTH. (B) MC2R and MRAP are expressed in brown adipocytes. Time course expression of MC2R and MRAP mRNA during brown adipocyte differentiation (n = 3). (C) Local BAT injection of ACTH increases both BAT and body temperature in anesthetized mice. (D) The thermogenic effects of ACTH (1 mg/kg), norepinephrine (NE, 1 mg/kg), and PBS in mice kept at room temperature. After determining the basal metabolic rate (BMR) at 30 C, respiration of mice treated with either ACTH (1 mg/kg), NE (1 mg/kg) or equal volume PBS via intraperitoneal injection (i.p.) was measured by indirect calorimetry for 1 hr at 27ºC. BMR and energy expenditure upon stimulation were quantified as area under the curve (AUC) (n = 6). Data are presented as means ± SD or individual values. (A) unpaired t test, (D) Two-way ANOVA (Tukey’s test).* = p < 0.05, ** = p < 0.01, *** = p < 0.001.

Taken together, multiple lines of evidence consistently demonstrated that ACTH harbors thermogenic activation capacity via directly acting on brown adipocytes, indicating that a rise in ACTH levels *in vivo* may contribute to BAT activation.

### Inhibition of MC2R signaling abolishes thermogenic effect of ACTH *in vitro* and impairs cold-induced energy expenditure *in vivo*

ACTH acts through the ACTH receptor complex, composed of the Gs protein-coupled receptor (GPCR) melanocortin2 receptor (MC2R) and the MC2R accessory protein (MRAP) (Cooray et al., 2008; Metherell et al., 2005). Notably, both MC2R and MRAP are primarily expressed in the adrenal gland and adipose tissue (Fig.2A). Consistently, knockdown of either MC2R or MRAP in brown adipocytes blocked ACTH- induced lipolysis (Fig.2B) and activation of UCP1-mediated thermogenesis (Fig.2C). The mediating effect of MC2R on ACTH-evoked brown adipocyte thermogenesis was further confirmed by using the high potent and selective MC2R antagonist CRN04894 (Kim et al., 2024). Biogenetic analysis revealed that CRN04894 preincubation resulted in a dose-dependent suppression of ACTH stimulated thermogenesis (Fig.2D). Notably, CRN04894 is also active *in vivo* and is currently being evaluated in clinical trials for diseases caused by an excess of ACTH (Kim et al., 2024). After validating its *in vitro* potency, we next explored the *in vivo* efficacy of CRN04894 in blocking ACTH action. Notably, oral administration of CRN04894 (60 mg/kg) suppressed cold-induced energy expenditure (Fig.2E). Together, these data strongly support that cold-stimulated ACTH activates brown fat thermogenesis through MC2R signaling.

**Figure 2.**
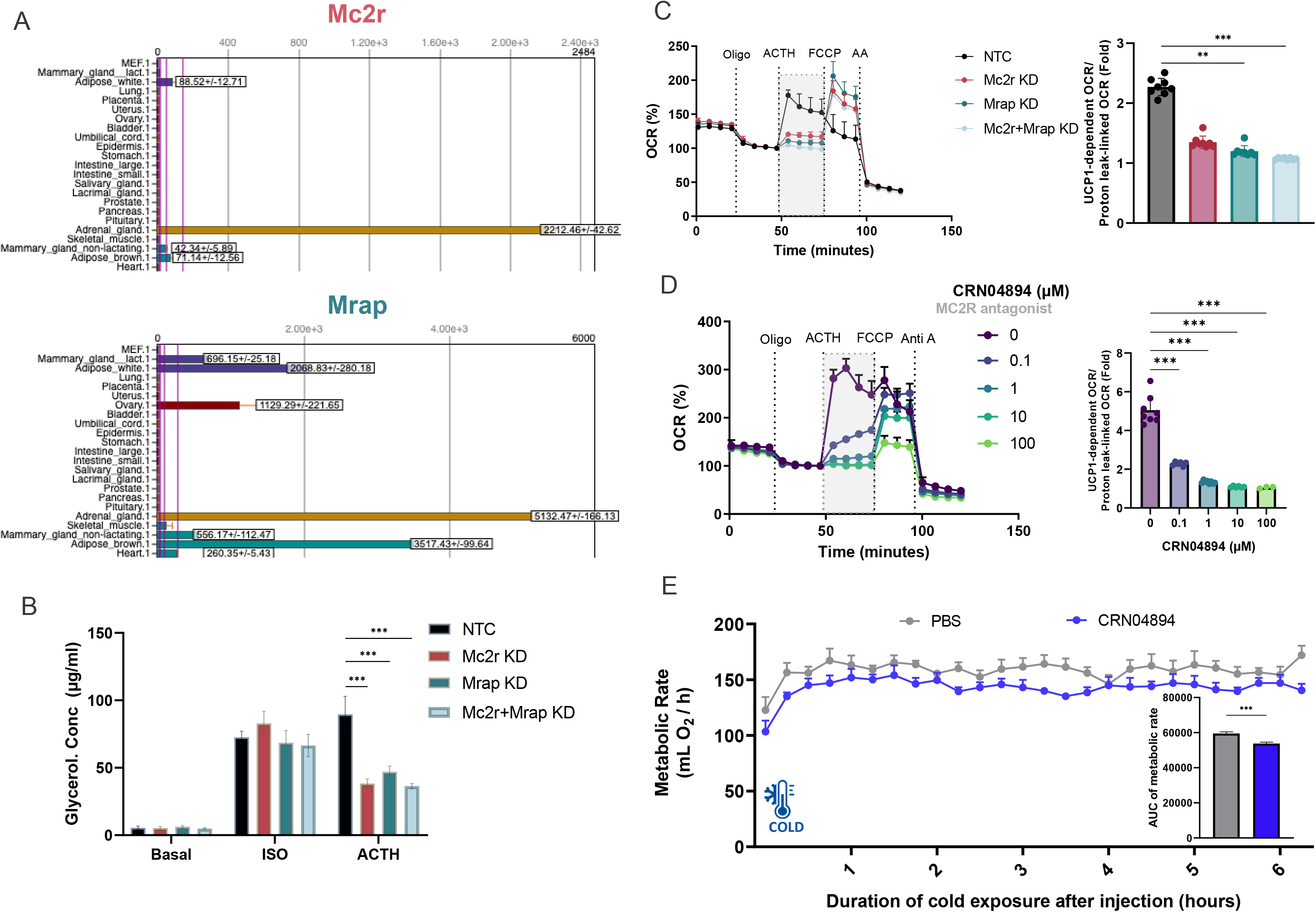
Inhibition of MC2R signaling abolishes thermogenic effect of ACTH *in vitro* and impairs cold-induced energy expenditure *in vivo*. (A) M2CR and MRAP are preferentially expressed in adrenal gland, brown and white adipose tissues according to the BioGPS database. (B) CRISPR-Cas9 based loss function of Mc2R and Mc2R-associated protein (MRAP) diminished the lipolytic effect of ACTH but not isoproterenol (ISO) in brown adipocytes. (C) CRISPR-Cas9 based loss function of both Mc2r and Mrap decreases the thermogenic effect of ACTH on UCP1-dependent respiration in brown adipocytes. UCP1-mediated uncoupled respiration (highlighted in grey) is expressed as fold increase of basal leak respiration after stimulation with ACTH (0.25 µM). (D) The MC2R antagonist CRN04894 blocks the ACTH-induced UCP1 activation in a dose-dependent manner *in vitro* in seahorse bioenergetics analysis. UCP1-mediated uncoupled respiration (highlighted in grey) is expressed as fold increase of basal leak respiration after stimulation with ACTH (0.25 µM). (E) Antagonism of MC2R with CRN04894 impaired cold-induced energy expenditure compared to vehicle control group in mice. Energy expenditure during cold exposure was quantified as area under the curve (AUC) (n = 6). Data are presented as means ± SD or individual values. (B) Two-way ANOVA (Tukey’s test), (C, D) One-way ANOVA, (E) unpaired t test.* = p < 0.05, ** = p < 0.01, *** = p < 0.001.

### Blocking the endogenous ACTH activity using a neutralizing antibody diminishes cold-induced energy expenditure and hyperphagia

Next, we assessed whether a physiological rise in ACTH levels upon HPA-axis activation is sufficient by itself to activate BAT thermogenesis *in vivo* under physiological conditions. Cold stress activated the HPA axis, as evidenced by a brief increase in the plasma levels of ACTH and a modest elevation in the level of corticosterone (Fig.3A). To block the endogenous ACTH activity during cold exposure, a neutralizing monoclonal ACTH antibody was applied. This antibody completely neutralized ACTH activity when tested *in vitro* (Fig.3B). Upon *in vivo* application, compared to vehicle controls, cold-induced increase in energy expenditure was largely attenuated in mice pretreated with the ACTH-neutralizing antibody ALD1613 (5 μg/mice) (Fig.3C). The effect size was comparable to ACTH antibody treatment (Fig.2E). Moreover, the release of glucocorticoids, downstream of HPA axis activation, was prevented, indicating a successful blockade of ACTH activity *in vivo* (Fig.3D). These results demonstrate that endogenous ACTH of the HPA axis contributes to cold-induced energy expenditure beyond its primary function in promoting the secretion of glucocorticoids (GCs) in the adrenal gland.

**Figure 3.**
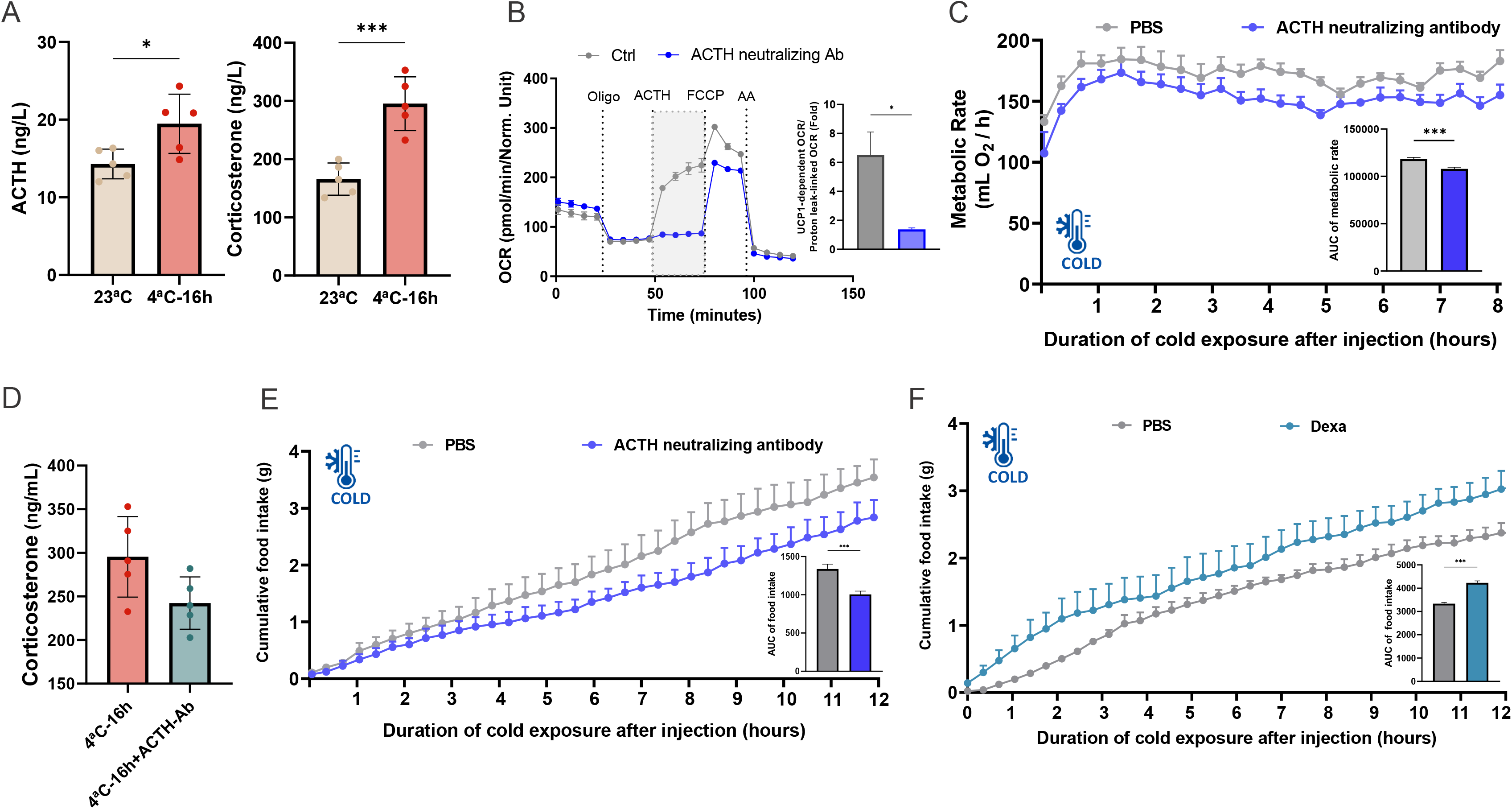
Blocking endogenous ACTH activity with a neutralizing antibody reduces both cold- induced energy expenditure and hyperphagia. (A) Cold exposure activates the HPA axis. Plasma ACTH concentrations and corticosterone levels after 16 h at 4 °C compared to 23ºC controls. (B) Blocking ACTH using a neutralizing antibody dampens UCP1-dependent respiration induced by ACTH. UCP1-mediated uncoupled respiration (highlighted in grey) is expressed as fold increase of basal leak respiration after stimulation with ACTH (0.25 µM). (C) Energy expenditure during cold-exposure in mice pretreated with ACTH neutralizing antibody and vehicle control. (D) Plasma corticosterone concentrations in mice treated with ACTH neutralizing antibody and vehicle control after 16 h at 4 °C. (E) Cumulative food intake of mice pre-treated with ACTH neutralizing antibody and vehicle control during cold-exposure. For statistical analysis, the area under the curve was calculated for cumulative food intake (see insert) (n = 6). (F) Glucocorticoids drive hyperphagia in mice. Cumulative food intake of C57BL6/J mice. Mice were injected (i.p.) with either dexamethasone (5mg/kg, Dexa) or vehicle (PBS) before feeding. For statistics, we calculated area under cumulative food intake time curve (see inset: AUC 0–12 hr) (n = 6) Data are presented as means ± SD or individual values. (A-F) unpaired t test.* = p < 0.05, ** = p < 0.01, *** = p < 0.001.

Cold stress not only rapidly activates energy expenditure, but also stimulates energy intake to compensate the energy deficit. Nevertheless, how energy expenditure and energy intake are coordinated to maintain energy homeostasis in the cold remains poorly understood. Unexpectedly, we found that upon neutralizing ACTH activity and therefore blocking the HPA axis, cold-induced hyperphagia was blunted as well (Fig.3E). Together, these data reveal that the HPA axis orchestrates energy expenditure and energy compensation during cold exposure.

### Glucocorticoids promote hyperphagia in mice

The HPA axis is a central neuroendocrine system composed of a cascade of hormonal signals. Upon activation, corticotropin-releasing factor stimulates ACTH release from the pituitary, which in turn triggers glucocorticoid secretion from the adrenal cortex. While glucocorticoids are best known for their gluconeogenic and anti-inflammatory actions in peripheral tissues, accumulating evidence shows that they also stimulate feeding behavior and bias food preference toward calorie-dense options (Dallman et al., 2004; Maniam & Morris, 2012; Sefton et al., 2016). We therefore hypothesized that glucocorticoid hormones, secreted as the end product of the HPA axis, mediate energy compensation during cold exposure. Consistent with this idea, glucocorticoid administration resulted in hyperphagia in mice during cold exposure (Fig.3F). For the underlying mechanism, we propose that glucocorticoids promote feeding by activating orexigenic neurons in the hypothalamus, such as agouti-related protein (AgRP) and neuropeptide Y (NPY) neurons. Supporting this model, AgRP and NPY neurons are known to express glucocorticoid receptor (GR) (Shibata et al., 2016); glucocorticoid response elements have been identified within the AgRP promoter region (Lee et al., 2013); glucocorticoids stimulate hypothalamic AgRP expression and activate AgRP neurons in rodents (Hagimoto et al., 2013; Perry et al., 2019; Shimizu et al., 2008); and cold-induced hyperphagia requires AgRP neuron activation in mice (Deem et al., 2020b; Yang et al., 2021). Collectively, these findings establish that HPA axis–derived glucocorticoids drive cold-induced hyperphagia, uncovering a physiological function that was previously unrecognized.

### Glucocorticoids have a permissive role on ACTH action

A physiologically important property of GPCRs is their tendency to desensitize upon exposure to their agonist (Freedman & Lefkowitz, 1996). To study desensitization in mature brown adipocytes, cells were pretreated with ACTH for 1 hour, washed and further cultured in differentiation media for 24 hours. The following day, cells were used for multiple assays to assess lipolytic function, cAMP production, or UCP1- dependent respiration (Fig.4A). ACTH-induced lipolysis was dampened in cells that had been pretreated the day before with ACTH compared to untreated cells (Ctrl) (Fig.4B). Interestingly, isoproterenol (ISO) or forskolin (FSK)-induced lipolysis was unaffected in cells pretreated with ACTH, indicating that the desensitization effect is only on ACTH signaling and not on cAMP production or the lipolytic pathway (Fig.4B). To our surprise, if cells were pretreated with dexamethasone (Dexa) for 30 minutes before the 1- hour pretreatment with ACTH, this synthetic glucocorticoid was able to rescue ACTH-dependent lipolysis the following day (Fig.4C). Dexamethasone did not only preserve responsiveness to subsequent ACTH stimulation in lipolysis, but also led to higher cAMP levels (Fig.4D) and UCP1-dependent respiration (Fig.4E). Dexamethasone rescued ACTH-induced MC2R desensitization in a dose-dependent manner (Fig.4F). Thus, beyond driving cold-induced hyperphagia, glucocorticoids also act permissively on ACTH by preventing MC2R desensitization, thereby maintaining cellular responsiveness to repeated stimulation. As ACTH–MC2R signaling governs both brown fat thermogenesis and rhythmic glucocorticoid release, this permissive effect underscores a broader principle of hormonal cross-talk in sustaining physiological adaptation.

**Figure 4.**
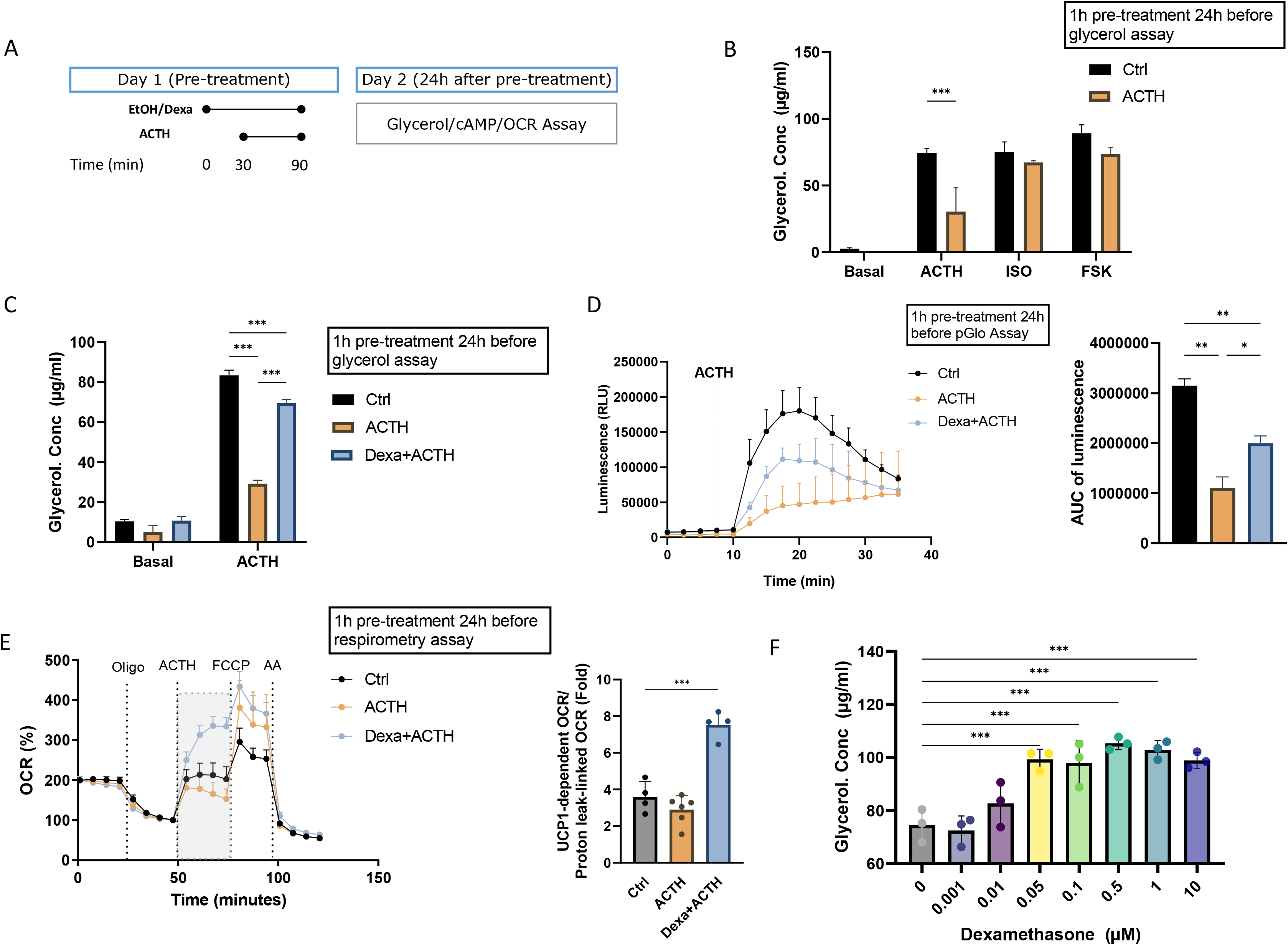
Glucocorticoids exert permissive effects on ACTH signaling through inhibition of ACTH- induced MC2R desensitization. (A) Experimental design to test the permissive effects of glucocorticoids on ACTH function. On Day 1, cells were pre-treated with dexamethasone (1 µM) for 30 minutes. Next, ACTH (0.25 µM) was added. 1 hour after the addition of ACTH, cells were washed and differentiation media was added. On Day 2, 24 hours after the pre-treatment, the assays of interest were performed such as glycerol, cAMP or oxygen consumption assays. (B) Pre-treatment of cells with ACTH leads to a lower response to ACTH, but not to FSK and ISO. Differentiated adipocytes were pre-treated with vehicle, ACTH, or ACTH+Dexa for on day one as explained in Figure 5A. On day 2, glycerol release-based lipolysis assay was performed under basal, ACTH (0.25 µM), ISO (1 µM), or FSK (10 µM) stimulated conditions. (C) Addition of dexamethasone during the pre-treatment preserves the responsiveness to of ACTH- induced lipolysis. Glycerol release in response to ACTH in brown adipocytes pretreated with ACTH ± dexamethasone. (D) Dexamethasone also preserves the cAMP response to ACTH in brown adipocytes pretreated with vehicle, ACTH, or ACTH+Dexa based on pGloSensor-22F reporter. (E) The permissive effects of dexamethasone are also observed in ACTH-induced UCP1-dependent respiration. Real-time bioenergetic profiling of brown adipocytes pretreated with vehicle, vehicle+Dexa, ACTH, or ACTH+Dexa by microplate-based respirometry (Seahorse XF96 Analyzer) under basal conditions and during successive addition of 5 µM oligomycin (oligo), 0.25 µM ACTH, 7.5 µM FCCP, and 5 µM antimycin A (Anti A). UCP1-mediated uncoupled respiration (highlighted in grey) is expressed as fold increase of basal leak respiration after stimulation with ACTH (0.25). (F) Dexamethasone exerts its permissive effects on ACTH-induced lipolysis in a dose-dependent manner. Glycerol release in response to ACTH in brown adipocytes pretreated with ACTH ± varying doses of dexamethasone. Data are presented as means ± SD or individual values. (B,C) Two-way ANOVA (Tukey’s test), (D-F) One-way ANOVA (Tukey’s test), .* = p < 0.05, ** = p < 0.01, *** = p < 0.001.

### Daily ACTH treatment elevates energy expenditure and promotes weight loss in diet-induced obese mice

Lastly, as our data demonstrates that ACTH can stimulate thermogenesis, ACTH treatment may hold promise for developing novel obesity therapies. To clarify the potential therapeutic usage, we daily injected diet-induced obese mice (DIO) mice with ACTH. ACTH treatment led to an increase in daily energy expenditure and a significant reduction in body weight (Fig. 5A-C). Furthermore, ACTH injections led to lower food intake in DIO mice (Fig. 5D). These data demonstrate ACTH also leads to thermogenesis in DIO mice. We conclude that the increase in energy expenditure and decrease in food intake after ACTH treatment strongly promotes a negative energy balance in DIO mice, presumably via the activation of brown fat.

**Figure 5.**
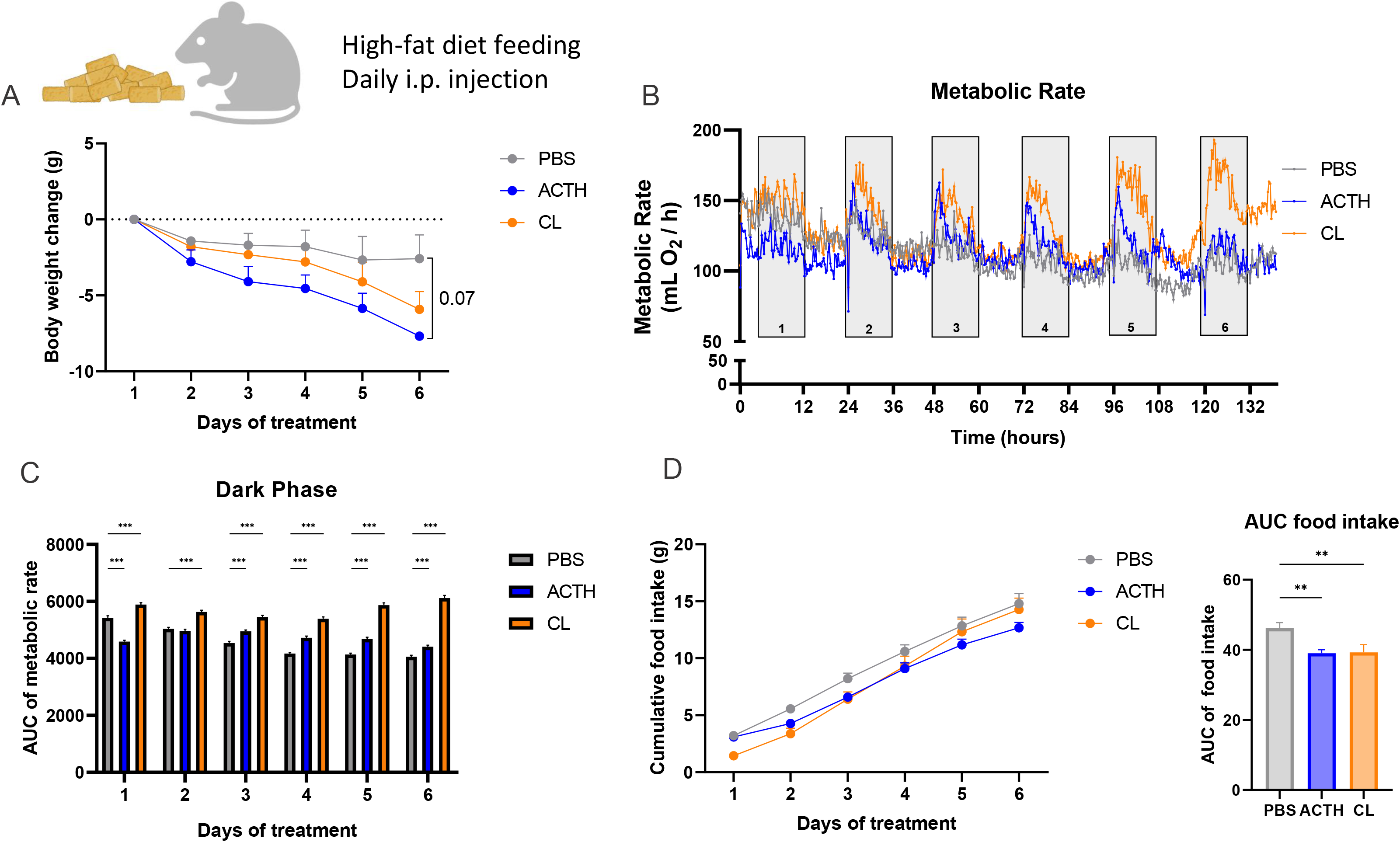
Daily ACTH injections increase energy expenditure and reduce body weight in diet-induced obese mice. (A) DIO male mice were injected daily with either ACTH (1 mg/kg ip), CL316,243 (CL; 1 mg/kg ip), or saline for 6 consecutive days. Daily body mass changes were calculated during the ACTH/CL316,243/saline administration. (B) Metabolic rate of DIO male mice injected with either ACTH (1 mg/kg ip), or CL316,243 (CL; 1 mg/kg ip) or saline for 6 consecutive days. (C) Area under the metabolic rate time curve was calculated for every day for the dark phase. (D) Cumulative food intake of DIO male mice were injected daily with either ACTH, CL, or saline. Data are presented as means ± SD or individual values. (A) analysis of covariance (ANCOVA) (C) Repeated-measures two-way ANOVA (Tukey’s test). (D) One-way ANOVA (Tukey’s test), .* = p < 0.05, ** = p < 0.01, *** = p < 0.001.

## Discussion

Energy expenditure and energy intake are coordinated to maintain energy homeostasis (Waterson & Horvath, 2015; Woods et al., 1998). To maintain such energy balance during cold exposure, the elevated energy demands imposed by thermogenesis must be offset by a corresponding increase in energy intake (Deem et al., 2020b). Significant knowledge gaps remain with respect to the homeostatic mechanisms involved in orchestrating energy balance during cold. It is vital to understand such intertwined coupling of energy expenditure and energy intake to define appropriate implementation strategies to manage the growing obesity epidemic, a consequence of an energy imbalance (Drenowatz, 2015). In this study, we uncovered an additional endocrine axis regulating brown fat thermogenesis and cold-induced energy compensation, whereby cold-induced ACTH, on one hand, in concert with norepinephrine released from sympathetic nerves directly activates BAT thermogenesis; one the other, stimulates glucocorticoid secretion to promote food intake to meet energy needs and compensate the energy deficit. We thus revealed an unappreciated physiological function of the HPA axis in orchestrating energy homeostasis via inter-organ communication during cold.

### The homeostatic mechanisms during cold exposure

Successfully surviving cold requires two simultaneous events. Firstly, generating substantial body heat by burning energy and secondly, acquiring sufficient energy to support heat production (Cannon & Nedergaard, 2004). Indeed, there is an innate biological drive toward coupling of energy intake and energy expenditure to maintain energy homeostasis in the cold. A deeper mechanistic understanding of how animals orchestrate energy intake and energy expenditure and maintain thermal and energy homeostasis is essential for understanding how organisms adapt to the continuously changing environment.

Upon cold exposure, a decrease in body core temperature and skin temperature eliciteds the primary cold thermoregulatory response, including cutaneous vasoconstriction and shivering thermogenesis (Morrison & Nakamura, 2011; Tan & Knight, 2018). Brown adipose tissue expressing uncoupling protein 1 (UCP1) is responsible for adaptive nonshivering thermogenesis giving eutherian mammals crucial advantage to survive in the cold. Classically, the thermogenic activity of brown fat is primarily driven by the SNS (Bamshad et al., 1999). However, novel factors capable of activating BAT through non-sympathetic mechanisms have been recently identified (Braun et al., 2018; Li et al., 2018, 2022). The identification of non-sympathetic controllers of BAT activity is of special biomedical interest as a prerequisite for developing pharmacological tools that influence BAT activity without the side effects of sympathomimetics (Braun et al., 2018). Endocrine circulating adrenocorticotropic hormone (ACTH) has long been demonstrated to harbor lipolytic and thermogenic activities (Boston & Cone, 1996; Cho et al., 2005; Hollenberg et al., 1961; White & Engel, 1958). However, the actual physiological significance and mechanisms of action remain incompletely defined. Through specifically blocking endogenous ACTH activity using a neutralizing antibody and preventing its action on brown adipocytes via an M2CR antagonist, we reveal that cold-induced endogenous ACTH in concert with norepinephrine released from sympathetic nerves directly activates BAT thermogenesis, which revealed an unappreciated physiological function of ACTH beyond its main role in regulating glucocorticoid secretion in the body. Notably, although an intact SNS-beta adrenergic signaling pathway is necessary for normal BAT morphology (Bachman et al., 2002), mice lacking beta-adrenoceptors (beta-less mice) upon cold- and subordination stress manifest restoration of UCP1 and BAT morphology, indicating the existence of alternative βAR- independent mechanisms of BAT activation (Razzoli et al., 2016). Even though whether ACTH could mediate the restoration of UCP1 and BAT morphology in mild-stressed beta-less mice has not been tested, the thermogenic capacity of endogenous ACTH demonstrated here suggests that ACTH may activate BAT under conditions of HPA axis activation beyond cold exposure. The physiological thermogenic activity of ACTH further indicates that neuronal and hormonal control exert overlapping yet synergistic regulation of brown adipose tissue function.

To maintain energy homeostasis during cold exposure, the increased energy demands of thermogenesis must be counterbalanced by increasing energy intake (Deem et al., 2020b). Classically, cold-induced hyperphagia is believed to be driven by the negative energy balance state that results from cold-induced thermogenesis, analogous to what occurs in other states of negative energy balance when body fat mass falls below a certain threshold. Recent studies demonstrated that the mere perception of cold rather than a negative energy balance is sufficient to elicit hyperphagia via increasing AgRP neuron activity (Deem et al., 2020b; Yang et al., 2021). In short, simply perceiving cold is enough to drive hyperphagia and thermogenesis, yet the mechanism by which cold information is conveyed to AgRP neurons is still unclear. Here, we demonstrat that, independent of a negative balance state, cold stress activates the HPA axis, and consequentially, HPA-axis-increased glucocorticoids serve as humoral signals and mediate cold-induced hyperphagia. Notably, AgRP neurons express glucocorticoid receptors, while glucocorticoids stimulate AgRP gene expression and neuron activity in rodents (Hagimoto et al., 2013; Perry et al., 2019; Shibata et al., 2016; Shimizu et al., 2008). Therefore, it is reasonable to propose that cold stress–induced increase in glucocorticoids via HPA axis activation stimulates AgRP neuron activity and promotes food intake to support energy homeostasis. Our findings thus reveal an endocrine system that is rapidly engaged during cold exposure and plays a critical role in driving the cold-associated hyperphagic response.

### Going beyond the classical function of the HPA axis

The primary function of the HPA axis is to orchestrate physiological adaptations to stress, which are essential for survival (Herman et al., 2016). In response to stress, neurons within the hypothalamic paraventricular nucleus (PVN) release the corticotropin-releasing factor (CRF) into the hypothalamo-hypophyseal portal system, thereby linking the hypothalamus with the anterior pituitary gland. CRF stimulates the secretion of ACTH from the anterior pituitary, which in turn binds to the melanocortin 2 receptor in the adrenal cortex to promote glucocorticoid (GC) release. Glucocorticoids exert their pleiotropic genomic actions through the glucocorticoid receptor (GR), a ubiquitously expressed transcription factor, which influences the stress response substantially. Through GC release, the HPA axis drives a range of adaptive responses to stressors, including mobilization of energy stores, enhancement of cardiovascular tone, and temporary suppression of non-essential processes such as growth, immune activity, and reproduction, while also restraining the stress response to prevent pathological overactivation (Sapolsky et al., 2000). More specifically, glucocorticoids not only elevate plasma catecholamine levels by inhibiting their extraneuronal uptake but also induce catecholamine supersensitivity in the heart by upregulating components of the β-adrenoceptor signaling cascade (Adameova et al., 2009). In addition, GCs exert permissive effects by enabling catecholamines to fully manifest their actions on cardiovascular tone (Hadcock & Malbon, 1988b).

In this study, going beyond the classical function of the HPA axis in the stress response, we reveal that cold stress elicits the activation of the HPA axis to confer protection at two distinct levels. Firstly, our study demonstrates that ACTH, released during stress, directly activates BAT thermogenesis. ACTH executes this thermogenic action by signaling via MC2R receptors abundantly expressed in brown adipocytes. This ACTH-driven thermogenic response is essential for acute cold adaptation, thereby assigning ACTH a previously unrecognized metabolic function as an endocrine activator of cold-induced BAT thermogenesis.

Secondly, we demonstrate that glucocorticoids, the end-products of the HPA axis, rapidly stimulate food intake to compensate for increased energy expenditure in the cold, thereby preventing excessive depletion of energy reserves while defending body temperature. Consistently, administration of GCs increases food consumption. Together, these findings uncover a novel molecular mechanism underlying cold-induced hyperphagia, positioning the HPA axis as a peripheral communication hub linking cold perception to central energy regulation. By revealing a non-canonical role of the HPA axis in orchestrating energy homeostasis beyond its traditional function in stress adaptation, our study highlights an unrecognized facet of the complex regulatory network supporting cold adaptation.

### The permissive effect of glucocorticoids on ACTH action

Glucocorticoids exhibit pleiotropic effects on animal physiology, ranging from permissive and suppressive to stimulatory and preparative actions, depending on context and target tissues (Munck & Náray-Fejes-Tóth, 1994; Sapolsky et al., 2000). While it is well established that glucocorticoids suppress ACTH secretion through a negative feedback, our study reveals an unappreciated permissive role of glucocorticoids in sustaining ACTH function. Like many G protein-coupled receptors (GPCR), MC2R undergoes desensitization and internalization following ACTH binding (Roy et al., 2011). Remarkably, glucocorticoids prevent ACTH-induced MC2R desensitization, thereby maintaining cellular responsiveness to repeated or sustained stimulation. Since ACTH-MC2R signaling regulates rhythmic glucocorticoid biogenesis and secretion, our findings hold broader physiological significance. The permissive action of glucocorticoids thus enables both ACTH-driven brown fat thermogenesis and rhythmic glucocorticoid release. In fact, the concept of permissive hormone action originated from early observations that changes in thyroid and glucocorticoid status dramatically altered cellular sensitivity to other hormones (Malbon et al., 1988). In adipose tissue, for example, glucocorticoids enhance catecholamine-induced lipolysis; adrenalectomy reduces norepinephrine-stimulated lipolysis, while glucocorticoid replacement restores it (Reshef & Shapiro, 1960; Shafrir et al., 1960; Shafrir & Steinberg, 1960). Essentially, glucocorticoids enable other hormones, such as catecholamines, to exert their full effects. Mechanistically, permissiveness is achieved via regulating hormone receptor expression, G proteins and adenylate cyclase activity (Allen & Beck, 1972; Lacasa et al., 1988). Our findings extend this concept by uncovering a novel facet of glucocorticoid action: regulation of GPCR desensitization. Traditionally, receptor activation and desensitization are considered tightly coupled, with ligands driving both signaling and subsequent attenuation, thereby constraining therapeutic potential. Here, we show that the permissive effect of glucocorticoids can uncouple these processes, a mechanism that may enable the design of more effective therapeutic strategies with broader efficacy and an expanded therapeutic window. Taken together, glucocorticoids exhibit dual roles: they suppress the HPA axis through a classical negative feedback while simultaneously exerting a permissive influence that allows ACTH to sustain brown fat thermogenesis and stimulate cortisol release during cold stress. These results highlight the importance of hormonal cross-talk in regulating energy balance and stress adaptation.

In summary, our results reveal an unappreciated physiological role of the HPA axis in coordinating energy homeostasis via inter-organ communication during cold exposure. Cold-induced ACTH, on one hand, in concert with norepinephrine released from sympathetic nerves directly activating BAT thermogenesis; one the other, stimulating glucocorticoids secretion to promote food intake to meet energy needs and compensate the energy deficit. Uncovering an endocrine axis regulating brown fat thermogenesis and cold-induced energy compensation not only advances our understanding of the thermoregulatory system, but also highlights potential strategies for obesity treatment by mitigating the accompanying hyperphagic response.

## Author Contributions

This study was designed by Y.L, M.L and D.W. Data were collected by M.L, X.G, Y.W, L.H, K.S and U.F. M.L analyzed the data. M.K., H.L., and H.U provided critical input throughout the project. The manuscript was written by Y.L and M.L with contributions from all authors. All authors read and approved the manuscript. M.L and X.G contributed equally to this study.

## Acknowledgments

This work was supported by grants from Forschungsgemeinschaft (DFG) [DFG-TRR 333/1 – 450149205 to Y.L, M.K, and H.U; Emmy Noether program (441904031) to Y.L; KL 973/20-1, #532683878 to M.K; SCHN 1696/1-1, #532683878 to K.S and FI 1546/8-1, #532683878 to U.F], European Research Council (ERC) [ERC starting grant: 101078516 to Y.L] and National Natural Science Foundation of China (NSFC) [32330012 to D.W].

